# Bone Marrow-Derived Cells Contribute to the Maintenance of Thymic Stroma including Thymic Epithelial Cells

**DOI:** 10.1101/2020.09.29.319426

**Authors:** Shami Chakrabarti, Mohammed Hoque, Nawshin Zara Jamil, Varan J Singh, Neelab Meer, Mark T. Pezzano

**Affiliations:** PhD Program in Biochemistry, The Graduate Center of the City University of New York; Department of Biology, City College of New York CUNY; PhD Program in Biology, The Graduate Center of the City University of New York

## Abstract

In paradox to critical functions for T-cell selection and self-tolerance, the thymus undergoes profound age-associated atrophy and loss of T-cell function, which are further enhanced by cancer therapies. Identification of thymic epithelial progenitor populations capable of forming functional thymic tissue will be critical in understanding thymic epithelial cell (TEC) ontogeny and designing strategies to reverse involution. We identified a new population of progenitor cells, present in both thymus and bone marrow (BM), that co-express the hematopoietic marker CD45 and the definitive thymic epithelial marker EpCAM and maintains the capacity to form functional thymic tissue. Confocal analysis and qRT-PCR of sorted cells from both BM and thymus confirmed co-expression of CD45 and EpCAM. Grafting of C57BL/6 fetal thymi under the kidney capsule of H2BGFP transgenic mice revealed that peripheral CD45+ EpCAM+ GFP-expressing cells migrate into the developing thymus and contribute to both TECs and FSP1-expressing stroma. Sorted BM-derived CD45+EpCAM+ cells contribute to reaggregate thymic organ cultures (RTOCs) and differentiate into keratin and FoxN1 expressing TECs, demonstrating that BM cells can contribute to the maintenance of TEC microenvironments previously thought to be derived solely from endoderm. BM-derived CD45+EpCAM+ cells represent a new source of progenitor cells that contribute to thymic homeostasis. Future studies will characterize the contribution of BM-derived CD45+EpCAM+ TEC progenitors to distinct functional TEC microenvironments in both the steady-state thymus and under conditions of demand. Cell therapies utilizing this population may prove useful for counteracting thymic involution in cancer patients.

## Introduction

The thymus serves as the primary lymphoid organ responsible for the development and selection of a self-tolerant T-cell repertoire(1, 2). Thymic epithelial cells (TECs) are the most significant component of the thymic microenvironment responsible for regulating T cell development and selection(2-4). Thus, proper organization and maintenance of TECs are critical for a properly functioning adaptive immune system(2, 3, 5). Despite the fundamental importance of the thymus in the development of T cells, the thymus undergoes profound atrophy relatively early in life(6-8). Thymus degeneration is physically evident in humans starting at puberty, where the loss of functional thymic microenvironments contributes to a progressively restricted naïve T cell repertoire, resulting in an immune system that is less capable of responding to new immune challenges(6, 9). Thymic involution also severely restricts the ability to generate long term, low morbidity tolerance to foreign transplants, including those of stem cell origin(10). Understanding the signals and cell-to-cell interactions that control both the development and postnatal maintenance of thymic epithelial cells is critical to design clinical strategies to counteract age-associated involution while enhancing thymic recovery following HSCT.

Structurally, the thymic stroma contains three compartments: 1) Cortex, 2) Medulla, and 3) Cortico-medullary junction(11). The cortex is composed of cortical thymic epithelial cells (cTEC), defined by primarily keratin 8 expression as well as the surface expression of BP1 and CD205, together with immature thymocyte subsets and it is involved in positive selection and MHC restriction during T-cell development(12). The thymic medulla is composed of medullary thymic epithelial cells (mTEC), defined by primarily Keratin 5/Keratin 14 expression together with UEA1 binding, and is involved in negative selection of mature single-positive (SP) thymocytes expressing autoreactive T-cell antigen receptors and development of FoxP3+Tregs(12, 13). The cortico-medullary junction is the perivascular space between the cortex and medulla, which serves as the entry point of CD4-CD8-(DN) thymocytes to the thymus(14) as well as the location where fully mature SP naïve T-cells exit to the peripheral blood vessels(2, 5, 12).

Most epithelial tissues are known to maintain tissue homeostasis throughout their adult life through the action of progenitor or stem cell populations(15). Previous work has shown that TECs of both the cortex and medulla develop from common progenitors during thymic organogenesis (16-19). The presence of progenitors restricted to each lineage has also been demonstrated(16, 20-24). Hamazaki et al., 2007 identified embryonic TEC progenitors, expressing high levels of claudin-3 and claudin-4, that exclusively give rise to mature mTECs(20). Progenitors initially expressing cTEC lineage markers(25), including the thymoproteosome subunit b5t(26, 27) and CD205(19), were shown to give rise to mTECS during thymic organogenesis. However, the mechanisms responsible for the maintenance of functional thymic microenvironments in the postnatal thymus remain unclear. Tissue maintenance might be mediated through stem-cell-based regeneration similar to tissues with high turnover rates, including hematopoietic, skin, and intestinal tissues(28). Alternatively, the thymus might be maintained by replication of more differentiated cells, similar to tissues with lower turnover rates, including the liver and pancreas(29). Gray et al. 2006 revealed that TECs, particularly mTECs, have much higher turnover rates than previously thought and are comparable with keratinocytes(30), implying a stem-cell-based regeneration of at least mTECs. Sekai et al. 2014 identified embryonic SSEA-1+ Cld3,4_hi_ mTEC stem cells that could maintain functional mTEC regeneration, including mature mTECs and central T cell tolerance(31). The clonogenic activity these SSEA-1+ Cld3,4hi TECs was rapidly decreased after birth in wild-type (WT) mice but was maintaining in postnatal Rag2 deficient mice, suggesting that crosstalk previously thought to be important in maintaining an appropriately organized thymus might inhibit maintenance of mTEC stem cell activity(31).

The existence of bipotent TEPC in the postnatal thymus has also been demonstrated(16, 23, 32, 33), and they appear to be found within the cTEC population(34)The stem cell characteristics of label retention and increased colony-forming capacity were identified within the Sca-1hi MHCIIlo TEC subset(32, 33). In contrast, Ulyanchenko et al. demonstrated bipotent TEPC potential within the PLET1+ MHCIIhi subset(23). These competing results suggest that more work is needed to identify the bipotent epithelial progenitor phenotype and the mechanisms responsible for postnatal TEC maintenance.

Hematopoietic stem cells (HSCs) give rise to a variety of hematopoietic cells found in the thymus, including thymocytes, B cells, dendritic cells, and macrophages. Several recent studies identified unexpected plasticity of bone marrow-derived hematopoietic stem cell populations(35-40). These studies suggested that bone marrow-derived HSC can give rise to a variety of different adult cell types. Krause et al. showed that BM-derived HSC when transplanted to irradiated hosts, home to and repopulate the bone marrow(35). They also showed that this cell population could migrate and differentiate into epithelial cells in the lung, GI tract, and skin(35). Theise et al showed, BM-derived HSC can give rise to hepatocytes in both rodents and humans(41, 42). However, these studies were refuted by showing that BM-derived populations begin expressing characteristics of non-hematopoietic populations as a result of cell fusion rather than by trans-differentiation in the hepatocyte system(43, 44). Later, Borue et al. used a Cre-lox system to demonstrate that even though a small subset of stem cells in the presence of somatic cells undergoes spontaneous fusion, there is a part of the HSC population that can undergo trans-differentiation to contribute to the keratinocyte expressing epithelial cells in the skin during the process of wound healing(45). They showed that BM-derived epithelial cells at the wound edges express Ki67, proving that these trans-differentiated cells are actively cycling(45). Wong et al. also showed that bone marrow-derived HSC could give rise to lung epithelial cells under conditions of naphthalene induced lung injury(40). Together, these studies showed unexpected plasticity, with BM-derived HSC giving rise to epithelial cells in different organs, but none of these studies suggested a contribution of BM-derived HSC to the maintenance of thymic epithelial or other thymic stromal components.

In this study, we describe a unique population of CD45+EpCAM+ cells in both adult BM and thymus that expresses both the definitive hematopoietic marker CD45 and the definitive thymic epithelial marker EpCAM. Using both organ transplant and reconstitution thymus organ culture, we show that this unique population of cells can contribute to the maintenance of non-hematopoietic components of thymic stroma, including keratin and FoxN1-expressing TECs.

## Materials and Methods

### Mice

C57BL6 and Actin H2BGFP and mRFP Rosa26 transgenic mice purchased from Jackson Laboratory were used for this study. The ages of the mice ranged from 7-9 weeks. All mice were bred and maintained at the City College of New York animal facility, and all experiments were performed with approval from the City College of New York institutional animal care and use committee.

### Antibodies

The following primary antibodies were used for experiments: CD45-PE Cy7, CD45-APC Cy7, CD45 APC (clone 30-F11, BD Bioscience), Pan-keratin (Polyclonal, Dako), EpCAM-PE (clone G8.8, eBioscience),FoxN1(clone G-20, Santa Cruz Biotechnology) FSP1(CloneS100A4, Biolegend),Thy1.2(clone 30-H12, BD Bioscience), CD11b (clone, BioLegend), CD11b PerCpcy5.5 (clone M1/70, Biolegend) CD4-biotinylated(clone RM4-5, BioLegend), CD8 – biotinylated (clone53-6.7, BioLegend), CD19 (clone1D3, Biolegend), Rabbit anti-GFP (Life),, and CD205 (LY75/DEC-205)(clone HD30 (Millipore)’We also used lineage depletion panel containing B220

The following secondary reagents were used for experiments: donkey anti-rabbit IgG-TRITC, donkey anti-rabbit IgG-Cy5, donkey anti-rabbit IgG-FITC, donkey anti-rat IgG-TRITC, donkey anti-goat IgG-FITC, goat anti-rat IgM-TRITC (Jackson ImmunoResearch), anti-rat IgG2a-FITC, anti-rat IgM-FITC, streptavidin-APC, streptavidin-APC Cy7, streptavidin-PerCP Cy5.5 (BD Bioscience) and streptavidin-TRITC (Southern Biotechnology Associate).

### TaqMan probes

All Taqman probes were purchased from Thermo-fisher Scientific, including CD45 (PTPRC), EpCAM, and 18S rRNA.

### Preparation of Thymic epithelial cells (TEC)

Thymic lobes were extracted from humanely euthanized mice in ice-cold phosphate-buffered saline (PBS). The thymic lobes were cleaned to remove fat and connective tissue and cut into four smaller pieces and transferred into a new tube containing PBS. The cut pieces were gently agitated using a glass Pasteur pipette to remove thymocytes. The remaining tissue was dissociated using Collagenase D (1.8mg/ml) and DNase I (1X) in HBSS solution incubated in a 37oC water bath for 15 minutes with gentle agitation every 5 minutes using a Pasture pipette. The single-cell suspension of digested tissue was washed with PBS and then filtered through a 100 μm strainer to remove any clumps of tissue. Hematopoietic cells were removed from the single-cell suspension using anti-mouse Thy1.2, CD11b, and CD19 antibodies and magnetic beads to enrich for the TEC population before sorting or flowcytometric analysis.

### Isolation of Bone Marrow cells

Bone Marrow cells were isolated from mouse hind limbs, as previously described(46). The resulting single-cell suspension was lineage depleted using biotinylated CD19, CD4, CD8, CD11b, CD11c, B220, Ter119 antibody, and biotin-binder magnetic beads. The post-depletion cell population was used for flow cytometric analysis.

### Flow cytometry

Cells were suspended in 100 μl of FACS staining buffer (FSB-5% fetal bovine serum, 5 mM EDTA in PBS) with appropriately diluted primary antibodies for 20 minutes on ice in the dark. After washing, secondary antibodies appropriately diluted in FSB were added, and the cells were incubated for an additional 20 minutes on ice in the dark. After washing, the cells were resuspended in 500 μl of FSB for data acquisition. Data acquisition was performed using an LSRII analyzer complete with three lasers (BD Bioscience), and cell sorts were performed using a FACS Aria (BD Bioscience). FACS data were analyzed using Flow Jo software (Tree Star) or FACS Diva software (BD Bioscience).

### RNA extraction and q-RT PCR from sorted cells

Cells were sorted as described above, and RNA was isolated using the Trizol method as directed by the Qiagen RNA isolation mini-prep kit. Isolated RNA was used for cDNA synthesis using Reverse transcriptase and random hexamers. The resulting cDNAs were used for q-RT PCR analysis using Taqman probes for CD45 and EpCAM. The 18s RNA housekeeping gene was used as a positive control to normalize the qPCR results.

### Reaggregate Thymic Organ Cultures (RTOC)

EpCAM+CD45+ cells were isolated and sorted to greater than 95% purity from the bone marrow, and thymus-derived from Actin H2BGFP mice and placed in single-cell suspensions as described above. The resulting highly purified EpCAM+ CD45+ GFP+ cells were mixed with dissociated fetal thymic stromal cells from an E14.5 C57BL6 fetal thymus at a ratio of 1:10 to form a reaggregate on a polycarbonate filter suspended over a trans-well plate. After 48 hours of culture, these reaggregates were then surgically transplanted under the kidney capsule of athymic Nude mice and grown for three weeks in vivo. After three weeks, the reaggregates were harvested and sectioned for further IHC analysis.

### Organ transfer experiments

Fetal thymi were isolated from E14.5 C57BL6 or Actin H2BGFP mice, as stated, and 5-6 fetal lobes were placed under the kidney capsule of either Actin H2BGFP mice or Rosa26 mRFP mice, respectively. Following a time course of 3, 6, 9, or 12 weeks the kidneys were isolated, and most of the exogenous thymi were used for flow cytometric analysis, while the remaining lobes were used for immunohistochemistry.

### Immunohistochemistry of Thymic Sections

Thymi were embedded in OCT compound from Tissue-Tek and snap-frozen using Liquid-N2. 8 mm sections were cut using a cryostat and mounted on coated slides. The slides were then fixed in 4% paraformaldehyde for 10 minutes, washed with PBS three times, then permeabilized using acetone for 10 minutes, washed and blocked with complete normal donkey serum (1% Normal donkey serum in Bovine serum albumin). Appropriately diluted antibodies specific to stromal subsets were added to the slides and then placed into a humid chamber at 4oC overnight. Secondary antibodies were applied, if needed, and incubated in a humid chamber for one hour at 37oC. After incubation, the slides were washed in PBS three times, mounted, then they were sealed following the application of Prolong Gold antifade +DAPI reagent. Images were acquired using a Zeiss LSM800 confocal microscope and LSM710 confocal microscope. Confocal images were analyzed using analyzed using LSM software (Zeiss).

### Statistical Analysis

Data comparisons were performed using a non-parametric, unpaired, one-tailed T-test. All graphs and statistical results were generated using Graph-pad Prism software. A P-value of <0.01 was considered significant.

## Results

Cells co-expressing both the definitive thymic epithelial marker EpCAM and the hematopoietic marker CD45 are present in both the bone marrow and the thymus.

### Presence of CD45+EpCAM+ cells in the thymus

Using standard enzymatic digestion of thymic tissue in conjunction with CD45 magnetic bead depletion to reduce the frequency of hematopoietic cells, we observed a persistent but rare population of cells that co-expressed both CD45+ and EpCAM+ (DP), shown in Figure 1A. This population has been described in several publications; however, they were identified as thymic nurse cells, which are complexes of cTECs with internalized CD45+ thymocytes(47). Due to the consistent presence of this DP population, even after very conservative doublet discrimination, we set out to determine if this unusual population truly represented thymic nurse cells or was a unique previously undescribed cell population expressing both CD45 and EpCAM. We enhanced the isolation of this population by depleting hematopoietic cells in our enzymatically dissociated thymic tissue using a cocktail of antibodies, including anti-Thy1.2, CD11b, CD11c, and CD19 together, rather than the usual an anti-CD45 antibody. Figure 1 shows a representative FACS profile demonstrating the increase in the frequency of the CD45+EpCAM+ population when anti-CD45 depletion was not utilized to deplete hematopoietic cells. Panel A shows that when anti-CD45 depletion was used to analyze the frequency of DP cells present in 8-month old C57BL/6 mice, the CD45+EpCAM+ population represents only an average of 0.23% (+/−0.2%) of the total cells present, whereas in panel B when a cocktail of anti-Thy1.2, CD11b, CD11c, and CD19 was used for depletion, the frequency of the CD45+EpCAM+ population in the total dissociated thymic cells increased to an average of 1.14%. (+/−0.3%). Every set of experiments had four replicates.

**Figure 1.**
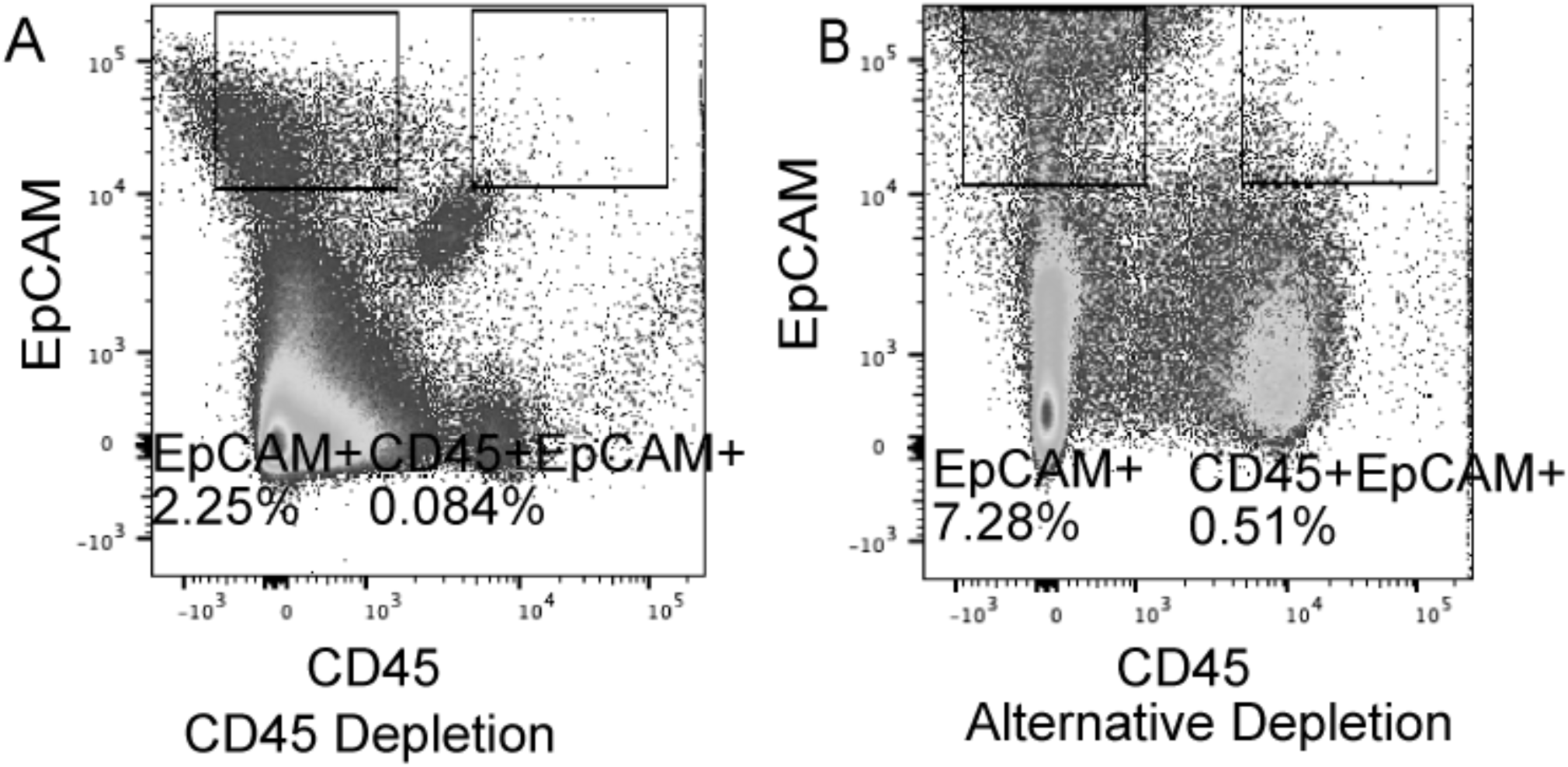
An alternative hematopoietic cell depletion method reveals a previously unidentified CD45+ EpCAM+ population in the thymus. FACS profiles show viable single cells gated on CD11c-, CD11b-and CD19- (lineage negative) cells. A- Shows the presence of only 0.084% of CD45+EpCAM+ cells present in dissociated thymic tissue when using sheep-anti-CD45 antibody followed by anti-sheep magnetic beads for depletion of hematopoietic cells. B-Shows enrichment of the CD45+EpCAM+ population to 0.51% derived from dissociated thymic tissue when a cocktail of sheep anti-Thy1.2, anti-CD11b and anti-CD19 antibodies followed by anti-sheep magnetic beads were used in place of anti-CD45. All the experiments were done in three replicates.

### Thymic CD45+EpCAM+ cells co-express both CD45 and EpCAM at both the protein and mRNA level

We reasoned that the apparent co-expression of CD45 and EpCAM, observed using flow cytometry of dissociated thymic tissue, could be the result of epitope sharing caused by the close association of thymocytes with thymic epithelial cells or due to penetration of CD45 antibodies into partially damaged TNC complexes. To confirm the presence of single cells co-expressing both CD45 and EpCAM, CD45+EpCAM+ cells were sorted to high purity (∼95% purity) using a FACS sorter from dissociated thymic tissue using conservative doublet discrimination. Cells were also gated to exclude hematopoietic cells using a cocktail of anti-Thy1.2, CD11b, CD11c, and CD19. To confirm that the DP cells expressed both the CD45 and EpCAM genes at the RNA level, RNA was isolated from CD45+EpCAM+; CD45+; EpCAM+; cell populations sorted from thymic tissue with greater than 95% purity and used for RNA isolation. All RNA samples were normalized to 5 ng/μl before performing qPCR. 18s rRNA was used as a housekeeping gene in order to compare relative amounts of RNA expression (Fig. 2B-C). Isolated and normalized RNA was analyzed using qRT-PCR with Taqman primers specific to EpCAM and CD45. Relative levels of RNA expression of both CD45 and EpCAM were normalized in each sorted population to the levels of 18s rRNA, to thoroughly rule out epitope sharing as the cause of this unique population. Figure 3 shows the fold change in the expression level for both CD45 and EpCAM between the two control populations (CD45+EpCAM-and CD45-EpCAM+) and the CD45+EpCAM+ population. Panel A shows the FACS representation of the sorted populations. Panel B shows the relative CD45 expression in all three populations, where the CD45+EpCAM-population serves as the positive control. As expected, the CD45+ control population has a high level of CD45 expression, whereas the EpCAM+ population has negligible (∼96-fold lower) CD45 expression compared to the CD45 only population. The DP population expressed ∼50 fold more CD45 than the CD45-EpCAM+ TEC population but had a ∼55-fold lower expression of CD45 than the CD45+ EpCAM-population. Panel C shows the relative EpCAM expression in all three populations, where the CD45-EpCAM+ population was used as a positive control. DP cells exhibited ∼52-fold higher EpCAM expression than the CD45+ population, while expression of EpCAM was ∼55% lower than that observed in the EpCAM only TEC positive control. As expected, the CD45 only population has negligible EpCAM expression (∼96 fold lower). The qRT-PCR confirmed that the DP population co-expressed both CD45 and EpCAM mRNA.

**Figure 2.**
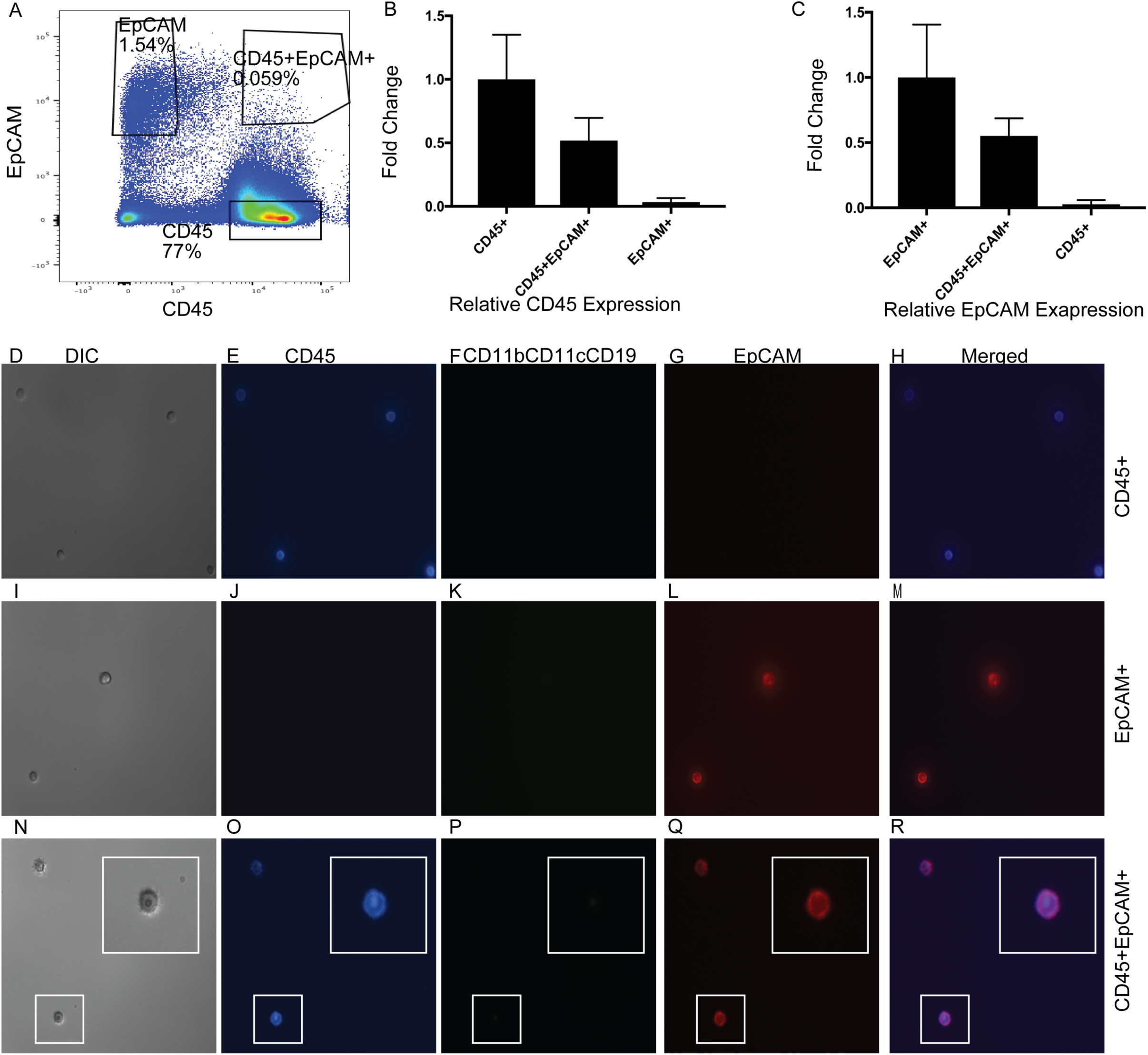
Single sorted thymic CD45+ EpCAM+ cells express both CD45 and EpCAM surface proteins and their mRNA. CD45+ EpCAM+ cells were sorted to high purity and analyzed for CD45 and EpCAM expression using immunofluorescence microscopy and qRT-PCR. CD45+EpCAM-; and CD45-EpCAM+ cells were sorted as controls. A: FACS analysis representation of gating strategy used for cell sort. B: Shows relative CD45 expression level in the thymus derived CD45+EpCAM+ cells compared to the CD45+EpCAM- and CD45-EpCAM+ control populations; C: Shows the relative EpCAM expression level in the thymus derived CD45+EpCAM+ cells compared to the CD45+EpCAM- and CD45-EpCAM+ control populations. D-H: DIC image, CD45 staining (blue), CD11b,CD11c,CD19 staining (green); EpCAM staining (red) and Merged image respectively for CD45+EpCAM-control population; I-M: DIC image, CD45 staining(blue), CD11b,CD11c,CD19 staining (green); EpCAM staining (red) and Merged image respectively for CD45-EpCAM+ control population; N-R: DIC image, CD45 staining (blue), CD11b,CD11c,CD19 staining (green); EpCAM staining (red) and Merged image respectively for the CD45+EpCAM+ experimental population. CD45+EpCAM+ cells of interest are enlarged in the insets to show colocalization of both proteins and represent the cells shown in the white boxes.

**Figure 3.**
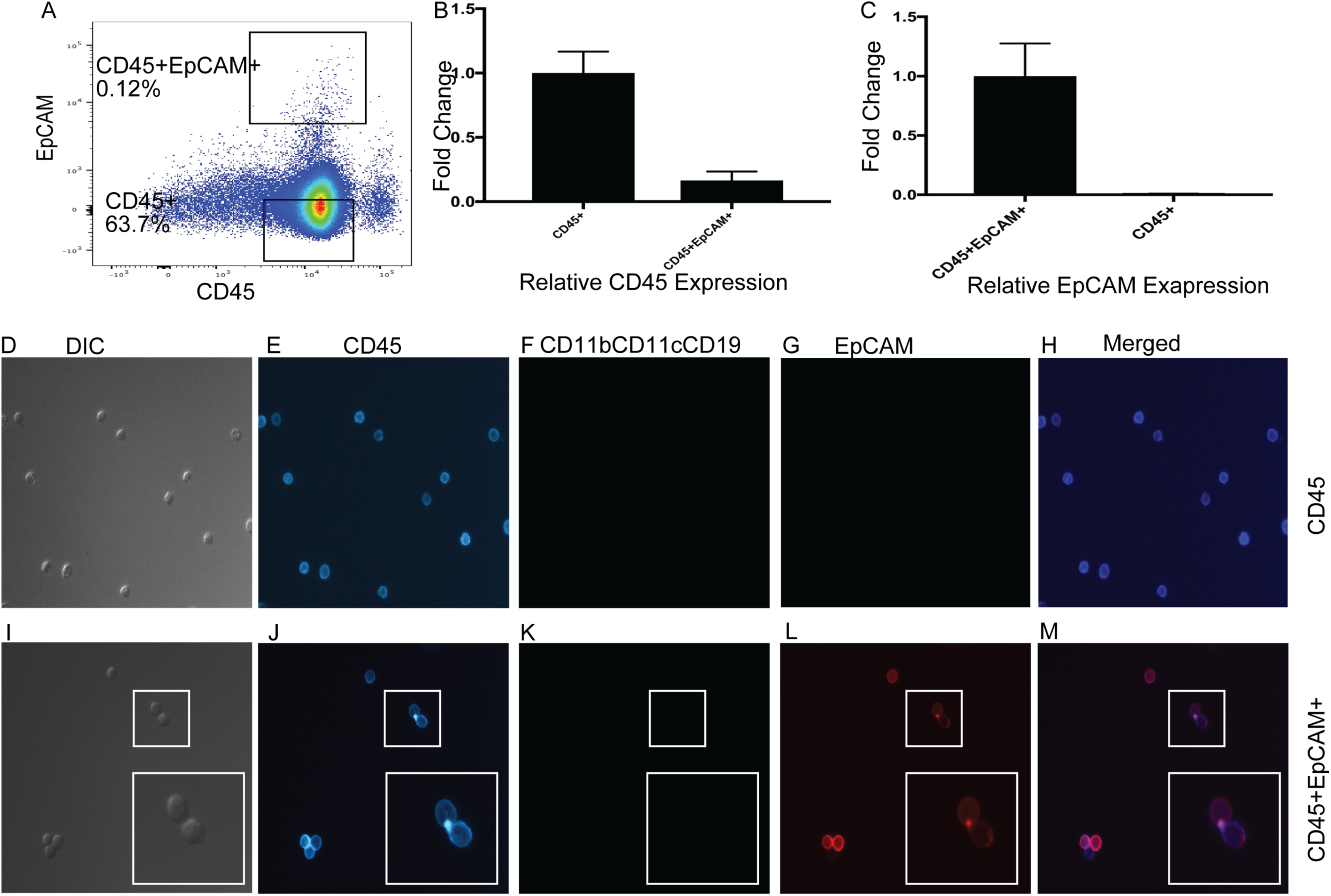
Single sorted bone marrow derived CD45+ EpCAM+ cells express both the CD45 and EpCAM genes and surface proteins. CD45+ EpCAM+ cells were sorted to high purity and analyzed for CD45 and EpCAM expression using immunofluorescence microscopy as well as qRT-PCR. CD45+EpCAM-cells were also sorted as a control. A: FACS profile of the gating strategy used for sorting. B: Shows the relative CD45 expression level in the sorted BM derived CD45+EpCAM+ cells compared to the CD45+EpCAM-control population; C: Shows relative EpCAM expression level in the sorted BM derived CD45+EpCAM+ cells compared to the CD45+EpCAM-control population. D-H sorted CD45+EpCAM-controls (D) DIC image; (E) CD45 staining (blue); (F) CD11b, CD11c, CD19 staining (green); (G) EpCAM staining (red); (H) Merged. I-M shows immunohistochemistry results obtained with sorted CD45+EpCAM+ cells (I) DIC image; (J) CD45 staining (blue); (K) CD11b, CD11c, CD19 staining (green); (L) EpCAM staining (red) and (M) Merged image. CD45+EpCAM+ cells of interest are enlarged in the insets to show colocalization of both proteins and represent the cells shown in the white boxes. All the experiments were done in three replicates.

Expression of both the CD45 and EpCAM surface proteins by sorted CD45+EpCAM+ cells was verified using immunohistochemistry. Cells that were either CD45+ EpCAM-or CD45-EpCAM+ were also sorted as controls and viewed under a fluorescence microscope to ensure that the antibodies were specific and that the CD45+EpCAM+ cells observed using flow cytometry were indeed single cells co-expressing both CD45 and EpCAM surface proteins. Figure 2 (D-R) shows the Immunohistochemistry (IHC) results obtained using the sorted CD45+EpCAM+ population and the control CD45+ EpCAM-or CD45-EpCAM+ populations from C57BL6 thymus following Anti-CD45 antibody and anti-EpCAM antibody staining. Combined CD11b, CD11c, and CD19 staining was performed on all sorted populations to rule out Langerhans cells that are known to co-express EpCAM(48-50) as well as to rule out non-specific binding of the anti-CD45 and anti-EpCAM antibodies to F_C_ receptors on hematopoietic cells, leading to a false-positive result. The CD45+EpCAM-control cells showed staining with only the anti-CD45 antibody (Fig. 2 D-H). The CD45-EpCAM+ control cells only exhibited anti-EpCAM staining (Fig. 2 I-M). However, the CD45+EpCAM+ cells stained strongly with both the anti-CD45 and anti-EpCAM antibodies while no staining with a cocktail of anti-CD11b, CD11c, CD19 antibodies was observed (Fig.2N-R) and confirming that the staining observed was not the result of non-specific F_C_-receptor binding. The inset of the merged image of the sorted CD45+EpCAM+ population shows a single representative cell expressing both CD45 and EpCAM. IHC results of the sorted cells confirmed protein co-expression of both CD45 and EpCAM on single cells.

To provide further evidence for the presence of this unique population and to localize the cells within the thymus, we sought to determine if the DP population could be seen within sections of an intact adult mouse thymus. Within adult mouse thymus, CD45+ cells far outnumber and are close to TECs. To better visualize the rare DP cells, within the background of abundant CD45+ thymocytes, we performed intraperitoneal injections of dexamethasone on 6-week-old C57BL/6 mice to specifically reduce the number of double-positive thymocytes, the most abundant CD45+ cells in the thymus. Three days following the dexamethasone injections, when the number of thymocytes is most reduced, the mice were sacrificed, and their thymi were harvested, sectioned, and stained with antibodies specific to CD45 (hematopoietic) and Pan-Keratin (epithelial) to enable identification of thymic stromal cells co-expressing CD45 and EpCAM. Confocal imaging of the stained sections revealed rare cells that share both hematopoietic (CD45+) and epithelial (Pan Cytokeratin) characteristics in the intact thymus (supplemental Figure 1). Together the IHC and qRT-PCR results obtained with highly purified sorted cells from mouse thymus and IHC of sections of intact thymus confirmed that the CD45+EpCAM+ cell population observed in dissociated thymic preparations truly co-express CD45 and EpCAM at both the protein and mRNA levels. Further, these results suggest that we and others in the field may be losing a large percentage of EpCAM+ cells by ignoring this CD45+ EpCAM+ population, including potential TEC progenitors.

### Rare CD45+ EpCAM+ Cells are present in the bone marrow

Bone marrow-derived cells, including hematopoietic stem cells (HSCs), were previously reported as contributors of epithelial cells for epithelial organs, including lung(40), gut, and uterus through the process of trans-differentiation(35). Wong et al. showed that the BM-derived CD45+CCSP+ cell population contributes to the alveolar epithelial cell population (CCSP is a unique alveolar epithelial cell marker)(39, 40). As the DP population is expressing the hematopoietic marker CD45, we wanted to determine if the CD45+EpCAM+ cells observed in the thymus might be derived from a population in the BM. Bone marrow preparations were analyzed for the presence of CD45+ EpCAM+ cells using flow cytometry. Multiple experiments with C57BL6 bone marrow showed that bone marrow contains a low frequency (∼0.1%-0.2%) of CD45+EpCAM+ cells (Figure 3A). To confirm that the population of CD45+ EpCAM+ cells detected in flow cytometry analysis actual represented single cells co-expressing both the CD45 and EpCAM proteins on their surface, and also expressed in mRNA level,we performed the same set of experiments shown above for the thymic CD45+EpCAM+ cell population. CD45+EpCAM+ cells and CD45+EpCAM-cells (control) were sorted to high purity (∼95%) from a bone marrow preparation depleted of lineage-committed cells and stained with CD45 and EpCAM antibodies. The sorted populations were subsequently analyzed using immunohistochemistry and qRT-PCR for expression of CD45 and EpCAM for each of the populations (Figure 3). To rule out epitope sharing as the cause of the bone marrow-derived DP population, we determined the relative level of mRNA expression of both CD45 and EpCAM compared to the housekeeping gene, 18s rRNA. All RNA samples isolated for these experiments were diluted to 5 ng/μL, to exclude differences in RNA amounts and normalized to 18s rRNA. The CD45+EpCAM+ population was shown to express both CD45 and EpCAM mRNA using qRT-PCR (Figure 3B-C). Figure 3 panel B shows the relative CD45 expression in the CD45+EpCAM+ population when compared with the CD45+EpCAM-positive control population. As expected, the CD45+ population has a high level of CD45 mRNA expression. The DP population clearly expresses CD45 mRNA. However, it has a ∼83-fold lower expression level than the CD45+ EpCAM-control population. Panel C shows the relative EpCAM expression in comparison to the CD45+EpCAM-, negative control population. The CD45 only population has a ∼100-fold lower EpCAM mRNA expression than the DP population.

The sorted DP and CD45 only populations in the BM were also analyzed using immunohistochemistry for surface protein expression of CD45, EpCAM, and a mixture of CD11b, CD11c, CD19 for each of the populations (Figure 3D-M). CD11b, CD11c, and CD19 combined staining was done to rule out F_C_ receptor binding to the CD45 and EpCAM antibodies leading to a false-positive result. The control CD45+EpCAM-population, only exhibited staining with the anti-CD45 antibody (3D-H). In contrast, the CD45+EpCAM+ cells were stained with both the anti-CD45 (3J) and anti-EpCAM antibodies (3L) while no staining with the CD11b, CD11c, CD19 cocktail (3K) was observed. The inset shows a higher magnification image of the DP cells (outlined by the box) for every staining panel and demonstrates that the DP cells represent single cells expressing both CD45 and EpCAM. Together these IHC results of sorted cells confirmed that single CD45+EpCAM+ cells co-express both the CD45 and EpCAM proteins on their cell surface.

Collectively, these data show that there is a rare population of previously undescribed cells in both adult thymus and the bone marrow that co-express the hematopoietic marker CD45 and the definitive thymic epithelial marker EpCAM. Further, we have demonstrated that in both adult bone marrow and thymus, single cells express both the CD45 and EpCAM surface proteins and that highly purified sorted populations express mRNA for both genes confirming that this rare and unique population exists. Absence of TNC complexes ensures that contrary to previous reports when careful double discrimination is used, the DP cells are not TNCs, while the absence of CD11c, CD11b, and CD19 surface protein expression allows exclusion of F_C_ receptor binding to antibodies or identification of Langerhans cells which are known to express both CD11c and EpCAM(48).

### CD45+EpCAM+ cells are recruited to thymic stroma from the periphery and contribute to EpCAM+CD45-TECs

In order to understand if peripheral CD45+EpCAM+ cells can contribute to the non-hematopoietic populations of the thymic stroma, E14.5 C57BL/6 fetal thymic lobes were transplanted under the kidney capsule of Actin H2BGFP transgenic mice. The engrafted thymic lobes were then harvested at different time points from 3 to 12 weeks following transplant and analyzed for the presence of H2B-GFP-expressing cells that have migrated into and contributed to the growing fetal thymi using both flow cytometry and immunohistochemistry. As a control, E14.5 C57BL/6 fetal thymic lobes were transplanted under the kidney capsule of C57Bl6 mice and harvested after three weeks for flow cytometric analysis to set up negative gating for GFP expression.

Figure 4A shows representative FACS analysis of dissociated transplanted thymic lobes analyzed at different time points after transplant, Fig 4B shows the percentage of CD45+EpCAM+GFP+ cells migrating into the C57BL/6 GFP-negative thymus, and 4C shows the frequency of CD45-EpCAM+GFP+ TECs that are derived from peripheral sources. Bar graphs of the mean percentages of peripheral CD45+EpCAM+GFP+ and CD45-EpCAM+GFP+ populations found in the transplanted thymi are shown in Figure 4D-E.

**Figure 4.**
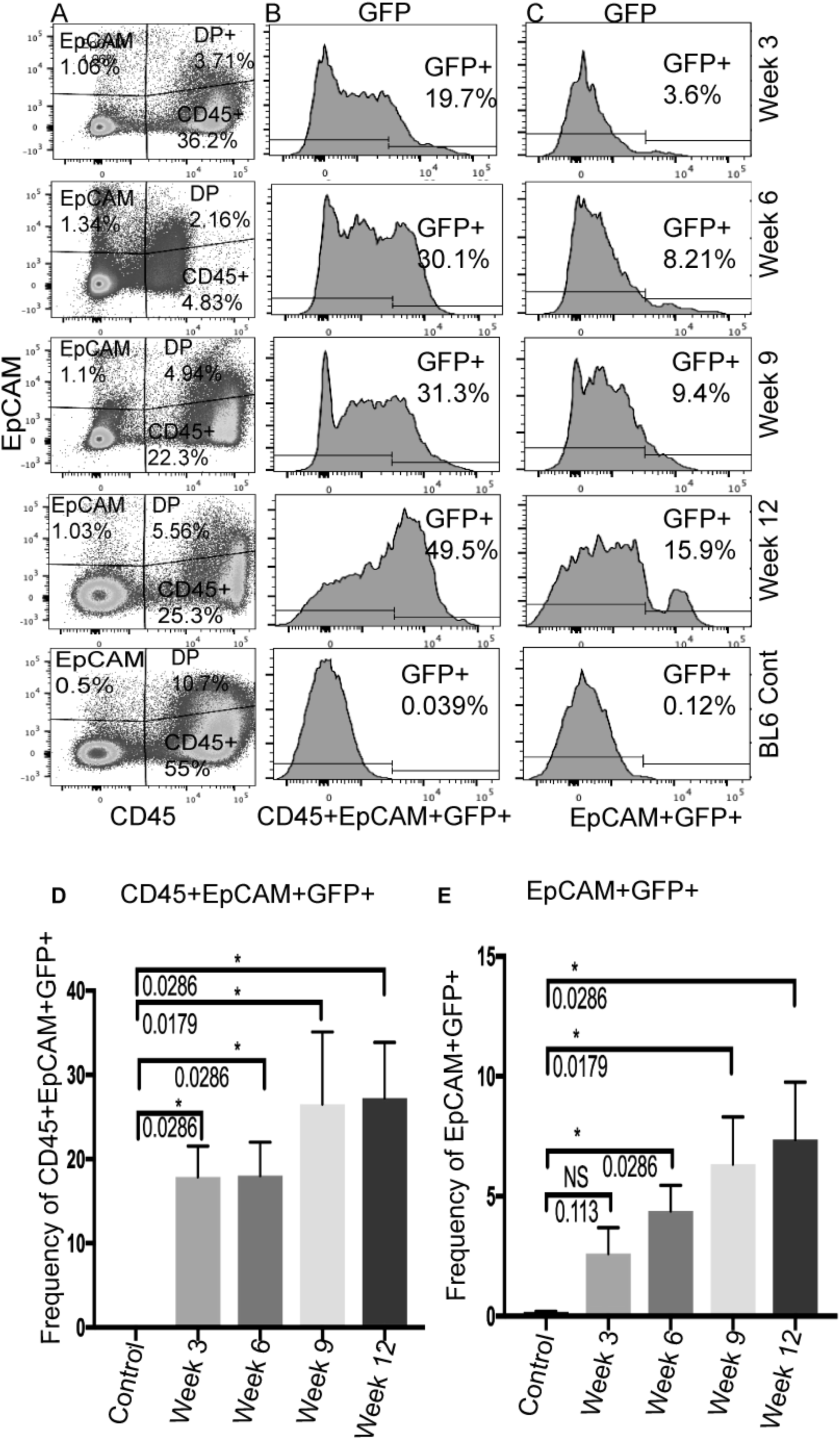
A peripheral population can contribute both CD45+ EpCAM+ cells as well as CD45-EpCAM+ TECs in the thymus. C57BL6 fetal thymi were transplanted under the kidney capsule of Actin H2BGFP mice and analyzed for the presence of GFP-expressing cells at 4 time-points 3 to 12 weeks after transplant using FACS. A: The FACS representation of the lineage depleted cells derived from dissociated engrafted thymi gated for CD45 and EpCAM. (B) The FACS representation of the percentage GFP+ cells within the CD45+EpCAM+subset found in the grafted thymus and (C) The FACS representation of the percentage GFP+ cells within the CD45-EpCAM+ population at the different timepoints after transplant under the kidney capsule. D. Bar graph showing the mean percentage of the GFP-expressing cells found within the CD45+EpCAM+ cell population at different time-point (error bars represent SEM). E: Bar graph showing the mean percentage of the GFP-expressing cells found within CD45-EpCAM+ TEC population at different time-points (error bars represent SEM). “*” indicates a statistically significant difference between control and any time-point. P-values for each t-test analysis are given with the bar graph. All the experiments were done in three-five replicates.

The flow-cytometric analysis revealed an influx of GFP+CD45+EpCAM+ cells into the engrafted fetal thymic lobes. The presence of CD45-EpCAM+GFP+ TECs was detected at a later time-point in the BL6 engrafts (Figure 4). An influx of the peripherally derived CD45+EpCAM+GFP+ population was observed at every timepoint. The average frequency of the CD45+EpCAM+GFP+ population remained almost constant at week three, 17.88% (+/− 7.3%), and week six, 18% (+/7.9%). At week nine and week twelve, the average frequency of the CD45+EpCAM+GFP+ population increased to 26.53% (+/−19%) and 27.3%(+/−13%), respectively. An unpaired, non-parametric one-tailed T-test demonstrated that there was a significant increase in the frequency of peripherally derived GFP+CD45+EpCAM+ cells within the C57BL6 fetal thymi that were transplanted under C57BL6 kidney capsules when compared with C57BL/6 control thymi transplanted under the kidney capsules of non-GFP expressing mice.

We compared each time point with the control. All the time-points analyzed, when C57BL/6 fetal lobes, where transplanted under the kidney capsule of ActinH2BGFP mice, showed a statistically significant number of CD45+EpCAM+GFP+ cells had migrated to the transplanted thymi.

Interestingly, a small but progressive increase in the frequency of CD45-EpCAM+GFP+ TECs was also observed, suggesting that true EpCAM-expressing TECs were also derived from peripheral sources. The average frequency of GFP expressing TECs increased from 2.6% (+/− 2%) at week three to 4.4% (+/− 2%) at week six. At week nine, the average frequency of GFP expressing TECs was 6.34% (+/− 4%) and increased to 7.4% (+/− 4%) at week twelve. C57BL/6 fetal thymi transplanted under the C57BL/6 kidney capsule were used as a control to allow gating on true GFP expressing cells in the engrafted thymus. When the frequency of EpCAM+GFP+ cells in the engrafted thymi at each time points was compared to the control (using a non-parametric, unpaired, one-tailed T-test), it showed significant numbers of EpCAM+GFP+ true TECs appearing in the transplanted thymic lobes. These results confirm that peripheral GFP+ CD45+EpCAM+ cells migrate into the thymus and that these peripheral cells then give rise to GFP-expressing CD45-EpCAM+ thymic epithelial cells. This result is unprecedented, as it suggests that contrary to previous models of thymic organogenesis, which suggested that all EpCAM expressing TECs were derived from the third pharyngeal pouch endoderm and its derivatives, that in fact up to 7% of EpCAM only TECs may be derived from peripheral sources that migrate into the thymus.

### Peripheral cells contribute to the pan-keratin and FoxN1 expressing epithelial cells in the thymic stroma

The engrafted thymi were also analyzed histologically in parallel experiments at the previously indicated time points. Figure 5 shows the staining pattern of PanK (an epithelial cell marker) and FoxN1 (a definitive thymic epithelial cell marker) in the transplanted thymic tissues at different time points following transplant. These histological results showed the presence of a PanK+ FoxN1+ H2B-GFP+ and PanK+FoxN1-H2BGFP population in the engrafted fetal thymus at week three and week six and PanK+FoxN1+H2B-GFP+ cells appeared in higher number at later time points (week nine and week twelve) shown in figure 5. Expression of PanK on a GFP+ cell in the transplanted thymi indicates that peripheral cells can migrate into the thymus and contribute to true TECs. At earlier time points, only PanK+FoxN1-GFP+ cells were observed, while at later time points, PanK+FoxN1+GFP+ cells were observed. This sequential emergence of two different populations of PanK+GFP+ cells that were initially FoxN1-followed by FoxN1+ cells with progressive age, indicates that first peripheral cells are recruited to the thymus and differentiate into only PanK expressing TEC, and at a later time point more mature PanK+FoxN1-expressing cells emerge.

**Figure 5:**
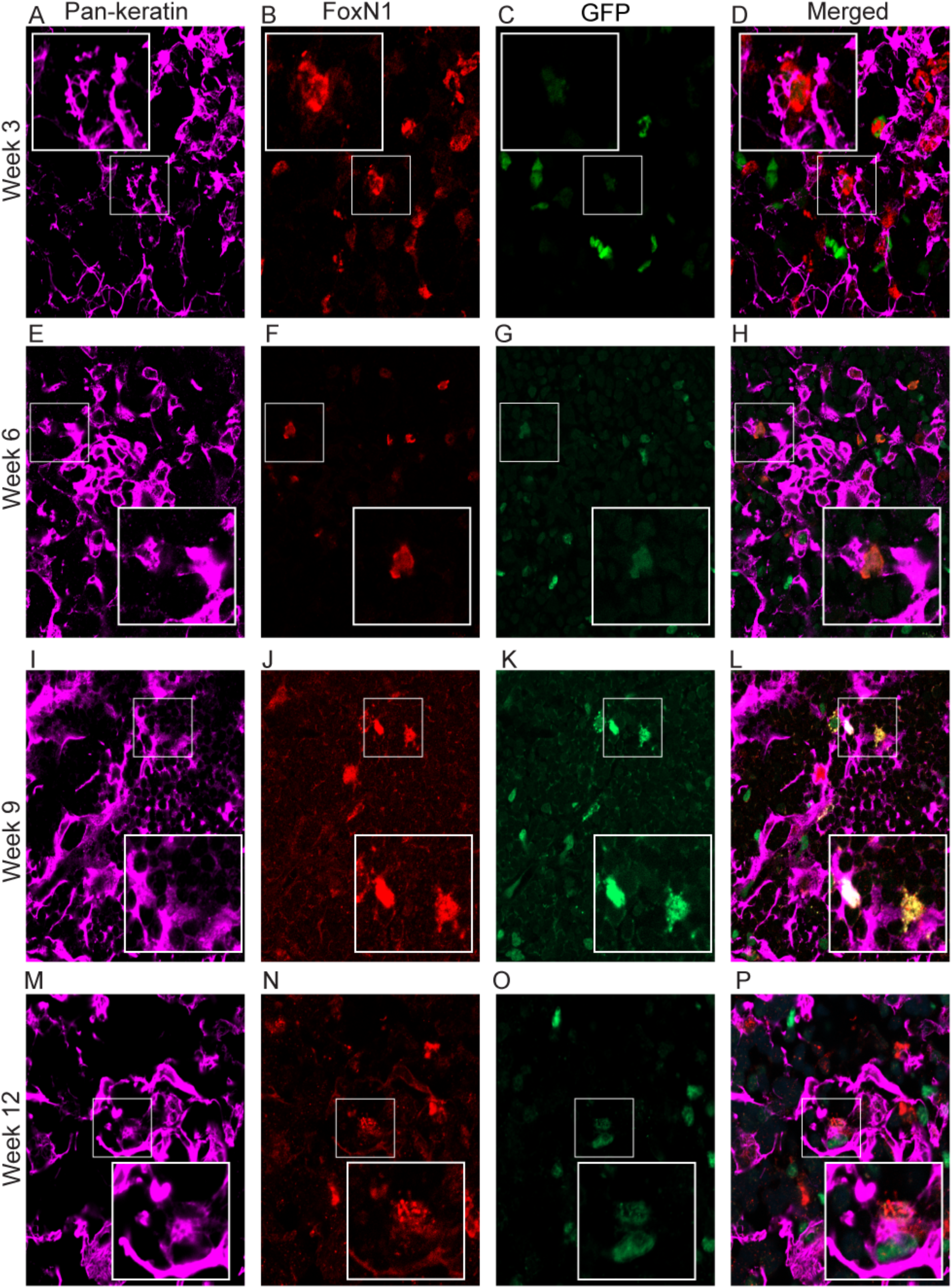
A peripheral population can migrate into the thymus and contribute to PanK and FoxN1 expressing thymic epithelial cells. C57BL6 fetal thymi were transplanted under the kidney capsule of Actin H2BGFP mice and analyzed for the presence of GFP-expressing peripheral cells 3-12 weeks after transplant using IHC. A-D: Week 3 transplanted thymic tissue sections showing expression of PanK (A), GFP+ peripheral cells (B), FoxN1 (C) and Merged image (D). E-H: Week 6 transplanted thymic tissue section showing expression of PanK (E), GFP+ peripheral cells (F), FoxN1 (G) and Merged image (H). I-L: Week 9 transplanted thymic tissue section showing expression of PanK (I), GFP+ peripheral cells (J), FoxN1 (K) and Merged image (L). M-P: Week 12 transplanted thymic tissue section showing expression of PanK (M), GFP+ peripheral cells (N), FoxN1 (O) and Merged image (P). GFP, Pank and FoxN1 expressing cells of interest are enlarged in the insets to show colocalization of both proteins and represent the cells shown in the white boxes. PanK+ FoxN1+ GFP+ cells and PanK+ FoxN1-GFP+ cells at all time points are indicated by white and red arrows respectively. All the experiments were done in three-five replicates.

### CD45+EpCAM+ cells can contribute to the thymic stroma in vivo

To understand the differential potential of the CD45+EpCAM+ population in vivo, we used reaggregate thymic organ culture (RTOC). GFP+CD45+EpCAM+cells were sorted to high purity (greater than 95%) from Actin H2BGFP mouse bone marrow or thymus and reaggregated with dissociated fetal thymic cells (E14.5) derived from GFP-C57BL6 mice. Reaggregates were cultured for 48hrs on polycarbonate filters supported by trans-well plates and then engrafted under the kidney capsule of athymic nude mice. The resulting ectopic thymic graft incorporated the GFP+CD45+EpCAM+ population into the thymic stroma. At week three, we observed PanK and FOXN1 expressing GFP+ cells in thymic stroma (Figure 6). RTOCs derived from bone marrow had more GFP+ TECs, shown in figure 6A-D, whereas thymus-derived RTOCs had a limited GFP+ TEC population (6 E-H). These results together showed that bone marrow and thymus derived CD45+ EpCAM+ cells could give rise to thymic epithelium and prove that peripheral cells contribute to the TEC component of the thymic stroma.

**Figure 6.**
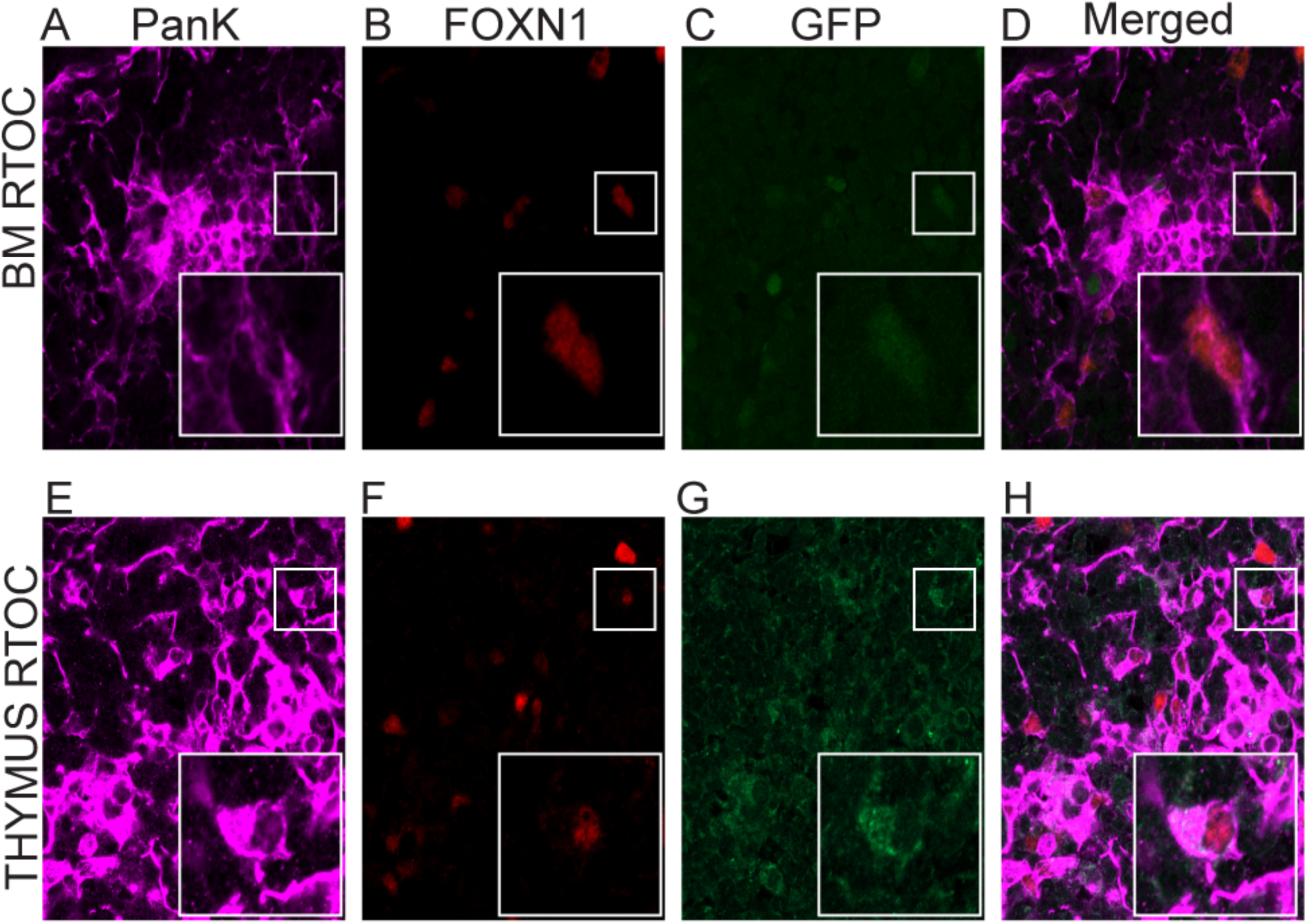
Reaggregate thymic organ cultures show that both bone marrow and thymus derived CD45+ EpCAM+ cells can give rise to PanK+ FoxN1+ thymic epithelial cells. (A-D) Sections of RTOC consisting of C57BL/6 fetal thymus mixed with sorted actin-H2BGFP-expressing bone marrow-derived CD45+EpCAM+ cells. A representative RTOC section showing PanK expression (A); FoxN1 expression (B) with GFP expressing cells derived from the CD45+EpCAM+ cells sorted from adult GFP-expressing bone-marrow (C), (D) Merged image of PanK, FoxN1 and GFP. (E-H) Sections of RTOC consisting of C57BL/6 fetal thymus mixed with sorted actin-H2BGFP-expressing thymic derived CD45+EpCAM+ cells. A representative RTOC section showing PanK expression (E); FoxN1 expression (F) and; GFP expressing cells derived from actin H2B-GFP-expressing CD45+EpCAM+ cells sorted from adult thymus (G); Merged image of PanK, FoxN1 and GFP (H). GFP, Pank and FOXN1 expressing cells of interest are enlarged in the insets to show co-expression of both proteins and represent the cells shown in the white boxes. All the experiments were done in three replicates.

### FSP1 expressing fibroblast cells in the thymus are derived from peripheral cells

During our FACS and IHC analysis, we observed that apart from the CD45+GFP+, CD45+EpCAM+GFP+, and CD45-EpCAM+GFP+ populations, there were a significant number of GFP+ cells that did not express either CD45 or EpCAM. In addition to TECs, another non-hematopoietic contributor of the thymic stroma is fibroblasts. Recently, Fibroblast Specific Protein-1 (FSP-1) expressing cells were shown to help to maintain medullary thymic epithelial cells(51). Engrafted thymi were analyzed for FSP1 expression using IHC (Figure 7). Figure 7A-P shows a representative figure of FSP1 staining in the transplanted thymi at different time-points. The frequency of peripherally derived GFP+ cells found in the transplanted fetal lobes that express FSP1 was calculated at the different time-points post-transplant. At week three, the percentage of GFP+ cells expressing FSP1 was 35% (+/− 8%), while at week six, it increased to 53 (+/−13). At week nine and week twelve, the percentage increased to 71.2% (+/−8%) and 71.4% (+/−10%), respectively(Supplemental figure 2). These results suggest that FSP1-expressing cells in the thymic stroma are continuously derived from cells migrating into the thymus from peripheral sources rather than through the expansion of stromal cells derived during embryonic development of the thymus.

**Figure 7:**
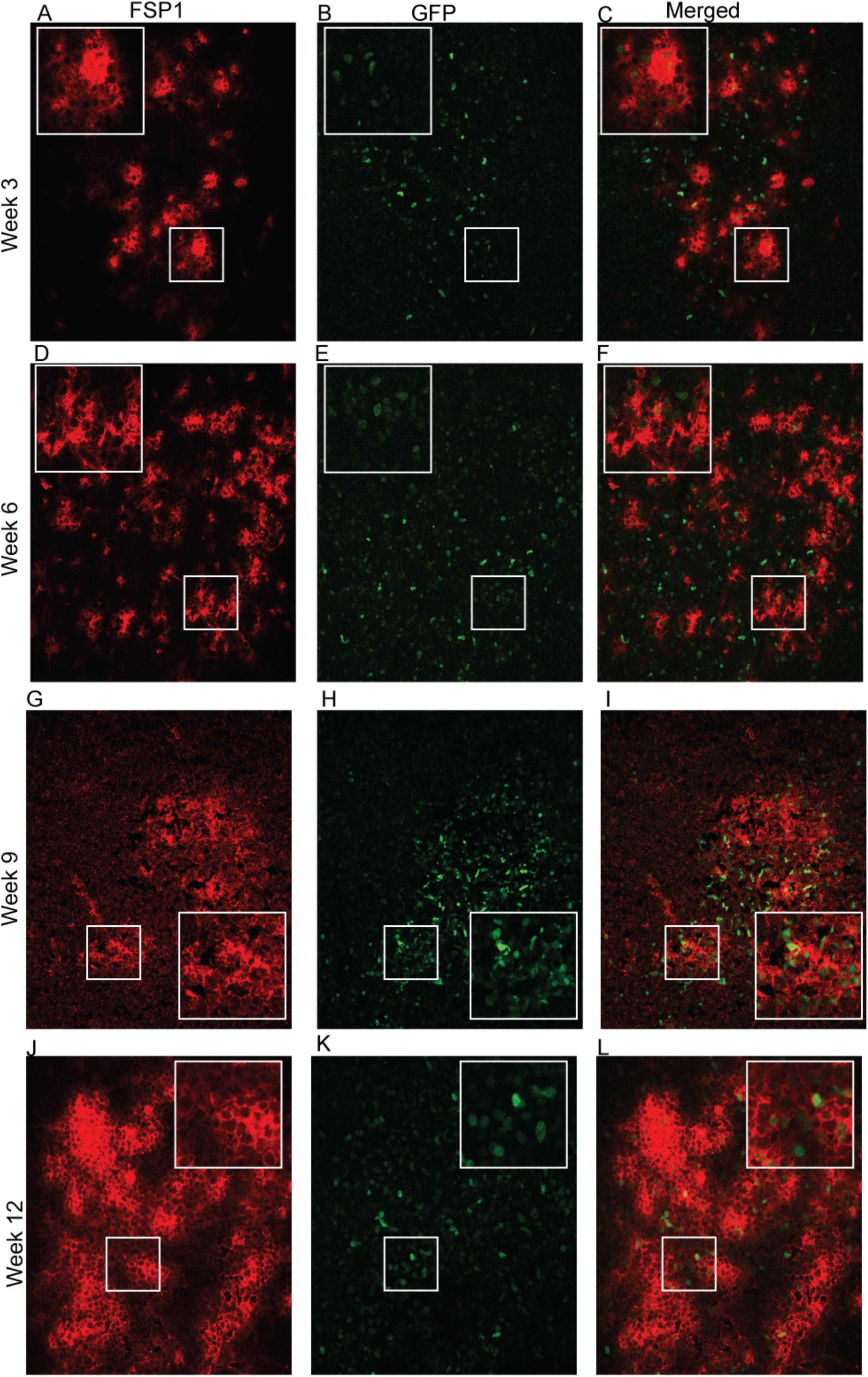
Peripheral cells migrating into the thymus contribute to FSP1-expressing fibroblasts. C57BL6 fetal thymi were transplanted under the kidney capsule of Actin H2BGFP mice and analyzed 3-12 weeks after transplant using IHC. A-C: Week 3 transplanted thymic tissue section showing the presence of GFP expressing peripheral cells (A), FSP1-expressing cells (B), and Merged image for FSP1 and GFP expression (C). D-F: Week 6 transplanted thymic tissue section shows the presence of GFP-expressing peripheral cells (D), staining pattern for FSP1 (E), and Merged image for FSP1 and GFP expression (F). G-I: Week 9 transplanted thymic tissue section shows presence of GFP expressing peripheral cell (G), staining pattern for FSP1 (H), and Merged image for FSP1 and GFP expression (I). J-L: Week 12 transplanted thymic tissue section shows the presence of GFP-expressing peripheral cells (J), staining pattern for FSP1(K), and Merged image for FSP1 and GFP expression (L). GFP, FSP1 expressing cells of interest are enlarged in the insets to show co-expression of both proteins and represent the cells shown in the white boxes. All the experiments were done in three-five replicates.

### CD45+EpCAM+ cells can contribute to the FSP1 expressing fibroblasts in the thymic stroma in vivo

We next wanted to investigate whether the BM and thymic derived CD45+EpCAM+ cells that contribute to TECs might also contribute to the peripheral populations giving rise to FSP1-expressing stromal components in the thymus. RTOCS composed of sorted CD45+EpCAM+H2BGFP+ BM or thymic cells mixed with non-GFP expressing fetal thymic cells were sectioned and stained with FSP1 to investigate if CD45+EpCAM+ cells can contribute to the FSP1+ fibroblast population of the thymic stroma (Figure 8). We observed FSP1 expressing H2B-GFP+ cells in the thymic stroma of RTOCs derived from both bone marrow and thymic-derived CD45+EpCAM+ cells showing that CD45+EpCAM+ cells can also give rise to FSP1 expressing fibroblasts.

**Figure 8.**
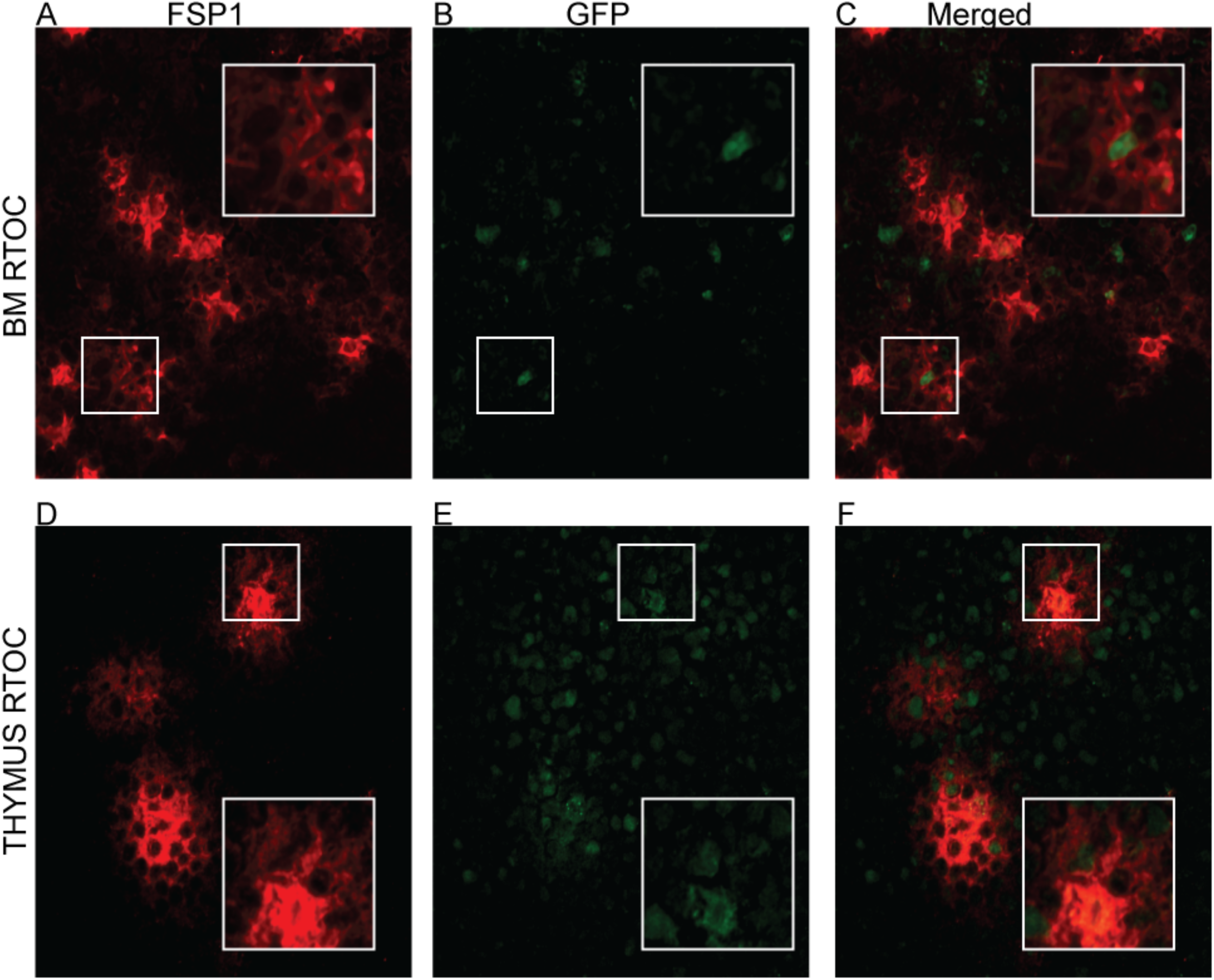
Reaggregate thymic organ cultures show that both bone marrow and thymus-derived CD45+ EpCAM+ cells can give rise to FSP1+ thymic stromal cells;(A-C) RTOC consisting of C57BL/6 fetal thymus mixed with bone marrow-derived actin H2B-GFP-expressing CD45+EpCAM+ cells. A representative RTOC section showing FSP1 expression (A); with GFP expressing cells derived from actin H2BGFP expressing CD45+EpCAM+ cells sorted from adult bone-marrow (B); Merged image of FSP1 and GFP (C). (D-F) RTOC consisting of C57BL/6 fetal thymus mixed with thymus-derived actin H2B-GFP-expressing CD45+EpCAM+ cells. A representative RTOC section showing FSP1 expression (D); with GFP expressing cells derived from CD45+EpCAM+ cells sorted from adult bone-marrow (E); Merged image of FSP1 and GFP (F). GFP and FSP1 expressing cells of interest are enlarged in the insets to show co-expression of both proteins and represent the cells shown in the white boxes. All the experiments were done in three replicates.

### Peripheral derived CD45+EpCAM+H2BGFP+ and CD45-EpCAM+H2BGFP+ cells are the results of trans-differentiation and not cell fusion

Previous studies reported that HSC cells could give rise to epithelial cells in different organs, including the liver, lung, GI tract, and skin through trans-differentiation(35, 40). However, these results were later disputed by Terada et al. and Wang et al., which suggested that BM-derived cells do not transdifferentiate, but they spontaneously fuse with other cells and adopt their characteristics(43, 44). The relatively high frequency of CD45+EpCAM+ cells observed in both the thymus and BM suggested that they were not derived from fusion. To determine if the H2BGFP-expressing TECs we observed in transplant experiments were the result of cell fusion events or true trans-differentiation, we transplanted E14.5 fetal thymuses derived from Actin H2BGFP time-pregnant mice under the kidney capsule of Rosa26 mRFP mice. If actin-H2BGFP+ EpCAM expressing CD45+EpCAM+ cells and the CD45-EpCAM+ true TECs shown in Figure 4 are the result of cell fusion, then in this double transgenic model both GFP and RFP should be expressed together on both the CD45+EpCAM+ cells and their resulting TEC progeny. However, if the cells are genuinely derived from peripheral sources, then co-expression of both RFP and GFP in the CD45+ EpCAM+ and EpCAM+ populations should not be observed. We analyzed the transplanted thymi at two different time points 6 and 9 weeks after transplant. Analysis of dissociated GFP-expressing fetal lobes using flow cytometry to identify RFP and GFP 6 and 9 weeks after transplant under the kidney capsule of Rosa26 mRFP mice revealed no co-expression of GFP and mRFP in either the CD45-EpCAM+, CD45+ EpCAM+ populations (Supplemental Figure 3.). mRFP+ peripheral cells were observed in all the populations, including CD45-EpCAM+ TECs. The CD45+EpCAM+ cell population had similar frequencies of peripheral mRFP+ cells at both time points; 68% (+/−15%) and 66% (+/− 15), respectively. The CD45-EpCAM+ population also contained ∼1% RFP + cells that were derived from peripheral mRFP cells migrating into the lobes. This set of results showed that peripheral CD45+EpCAM+ cells that migrate to the thymus and contribute to the thymic stroma are not the result of cell fusion and any H2BGFP+CD45+EpCAM+ cells observed as well as their H2BGFP+ CD45-EpCAM+ TEC progeny represent peripheral cells migrating into the thymus and contributing to stromal components.

## Discussion

In this study, we describe a unique, rare population in the postnatal bone marrow, which can migrate to the thymus and contribute to the thymic epithelial cell pool. This bone marrow-derived population expresses both the definitive thymic epithelial cell marker EpCAM and pan-hematopoietic cell marker CD45. This unique population was also identified in dissociated thymic tissue as well, supporting the idea that it migrates from the bone marrow to the thymus. Immunohistochemistry and qRT-PCR results performed with highly purified sorted populations of these CD45+EpCAM+ cells derived from both the bone marrow and thymus showed that these cells express both CD45 and EpCAM at both the mRNA and protein levels, confirming that they are not thymic nurse cells as previously reported(47) or an artifact derived from enzymatic dissociation of closely associated TECs and thymocytes (Figure 2 and Figure 3).

Several previous studies suggested that bone marrow-derived populations could contribute to the epithelial cell populations of different organs, including lung, stomach, intestine,uterus(35, 38, 40, 41); however, these studies never identified a contribution to the thymus. In this study, for the very first time, we are showing the presence of a potential thymic epithelial progenitor cell population within the bone marrow. Analysis of C57BL6 fetal thymi, after being transplanted under the kidney capsule of Actin H2BGFP mice, revealed that the CD45+EpCAM+ population can migrate from the periphery to a growing fetal thymus and can contribute to non-hematopoietic components of the thymic stroma (Figure 4). With progressive time after transplant, the frequency of the CD45+EpCAM+GFP+ cell population increased and was followed by the subsequent emergence of EpCAM+GFP+ TECs in the transplanted thymus.

IHC analysis of the same tissue showed the presence of Pan-keratin and GFP expressing cells in the transplanted tissues at all time points (Figure 5). Interestingly, Pan-keratin+FoxN1+GFP+ cells also appear to emerge at later time points. Foxn1 is a crucial transcription factor in TEC development and thymic function, and it is a well-established marker for thymic epithelial cells. The sequential expression of FoxN1 leads us to believe that peripheral cells can migrate to the growing thymus and can contribute to thymic stroma as PanK expressing thymic epithelial cells, which later differentiate into more mature FoxN1 expressing TEC at later time points(52, 53).

The presence of relatively abundant GFP+ cells in grafted fetal thymic lobes that did not bind the pan-keratin antibody, CD45, or Foxn1 led us to analyze the engrafted lobes for the presence of other non-hematopoietic stromal components including fibroblasts using FSP1 (a fibroblast marker). Recently it has been shown that FSP1 expressing cells are essential for the maintenance of medullary thymic epithelial cells(51). FSP1 staining of the transplanted tissue revealed the presence of a GFP expressing peripherally derived FSP1+ cell population (Figure 7). We also observed with progressive time that the number of peripherally derived GFP+FSP1+ cells increased (Supplemental Figure 2).

These results together showed that a peripheral CD45+EpCAM+ cell population could migrate to the thymus and contribute to the non-hematopoietic components of the thymic stroma. The CD45+EpCAM+ population appears to contribute to thymic stroma in two different ways: 1) by directly giving rise to EpCAM and keratin expressing epithelial cells and 2) giving rise to an FSP1 expressing cell population, known to support the maintenance of the medullary thymic epithelial cell population. Limitations in the availability of antibodies to sort FSP1-expressing cells or co-localize both Keratin and FSP1 on the same populations post-RTOC precluded determining whether they represented the same population or unique lineages derived from the CD45+EpCAM+ population. Future experiments will need to be performed to determine the lineage relationship between FSP1+ GFP + peripheral cells and PanK+GFP+ peripherally derived TECs.

Although several previous studies showed that bone-marrow-derived populations including HSCs could contribute to tissue regeneration in epithelial organs, some of this work had been refuted by subsequent studies which demonstrated that bone marrow-derived cells undergo fusion allowing them to take on characteristics of the recipient organs cell phenotype, instead of genuinely transdifferentiating(43). Our results obtained when ActinH2BGFP fetal thymus was transplanted under the kidney capsule of mRFPRosa26 mice clearly showed that the CD45+EpCAM+ RFP+ cell population observed in the ActinH2BGFP thymus after transplant is not the result of fusion (supplemental Figure 3) since no cells co-expressing RFP and GFP were observed. The RFP+ CD45-EpCAM+ TECs observed in the transplanted thymi are derived from peripheral RFP+ cells migrating into the GFP-expressing fetal lobe.

When highly purified CD45+EpCAM+ GFP+ cells from the bone-marrow were reaggregated with non-GFP fetal thymic stroma, the GFP+ bone-marrow-derived cells gave rise to pan-keratin and FoxN1-expressing true thymic epithelial cells(Figure 6A-D), as well as FSP1 expressing fibroblast cells(Figure 8A-C). Similar results were obtained with reaggregates made using the thymus-derived CD45+EpCAM+ population (Figure 6E-H and Figure 8D-F). Together these results confirm that the CD45+EpCAM+ cell population from both bone marrow and thymus can differentiate to Pan-Keratin expressing epithelial cells as well as FSP1 expressing fibroblasts. Our results suggest that the thymus-derived CD45+EpCAM+ cells in RTOCs exhibit less potential to differentiate into of Pan-keratin expressing TECs than BM-derived CD45+EpCAM+ cells supported by the lower frequency of Pan-Keratin+ true TECs. This result leads us to hypothesize that the BM-derived cells have more potential to differentiate in Pan-Keratin expressing TECs, whereas the thymus-derived CD45+EpCAM+ population might have lost some of its potentials or some of the cells may be committed to a different fate other than Pan-Keratin expressing epithelial cells.

In summary, this study demonstrates that a bone-marrow-derived population expressing both CD45 and EpCAM can migrate to the thymus and can give rise to two different non-hematopoietic cell populations in thymic stroma including both FSP1-expressing fibroblasts and keratin/FoxN1-expressing TECs. It is not clear whether the CD45+EpCAM+ cells are derived from HSCs or represent a distinct lineage of BM cells. Future studies will also need to be performed to determine the lineage relationships between the FSP1+ cells and the true keratin expressing TECs derived from the CD45+EpCAM+ population. They could represent distinct fates of the same population or steps in an undefined ontogeny. It is also unclear how significant the CD45+EpCAM+ population and its progeny are to TEC homeostasis or how they might contribute to self-tolerance or T cell selection. Previous work has suggested that the MHC background of the thymic microenvironment determines the MHC restriction of developing T cells after BMTs(54). If BM-derived cells are migrating into the thymus and contributing to the maintenance of TEC subsets, then it is surprising that they do not influence the MHC restriction of developing thymocytes. Further studies needed to be done to understand the ontogeny of this population and its contribution to distinct TEC subsets. If migrating CD45+EpCAM+ cells only contribute to mTECs, then it is possible that they do not influence MHC restriction but might contribute to self-tolerance through negative selection or the development of Regulatory T-Cell (T_regs_).

Despite the presence of a progenitor cell population in BM that can contribute to TECs, the thymus still undergoes age-related involution. Understanding the lineage of the CD45+EpCAM+ BM population as well as the mechanisms responsible for the migration of this population to the thymus and how they contribute to the maintenance of TEC number and organization will be critical in counteracting age-associated involution, particularly in cancer patients, due to enhanced degeneration in response to therapy. This study addresses a significant unmet clinical need for thymus reconstitution by defining the contribution of bone-marrow-derived thymic epithelial progenitor cells (TEPC) to the maintenance of TEC microenvironments in the postnatal thymus. It should provide an additional target for overcoming thymic atrophy and hence, the development of more strategic therapies for immunological-based diseases and cancer.

## Acknowledgements

This work was supported by NIH/NIAID 9SC1AI04994, NIH-NCI U54CA137788/U54CA132378, NIMHHD 8G12MD007603. We would like to thank Jeffrey Walker for his flow cytometry support.

