## Supplemental Figures S1-S3 for "Bone Marrow-Derived Cells Contribute to the Maintenance of Thymic Stroma including Thymic Epithelial Cells"

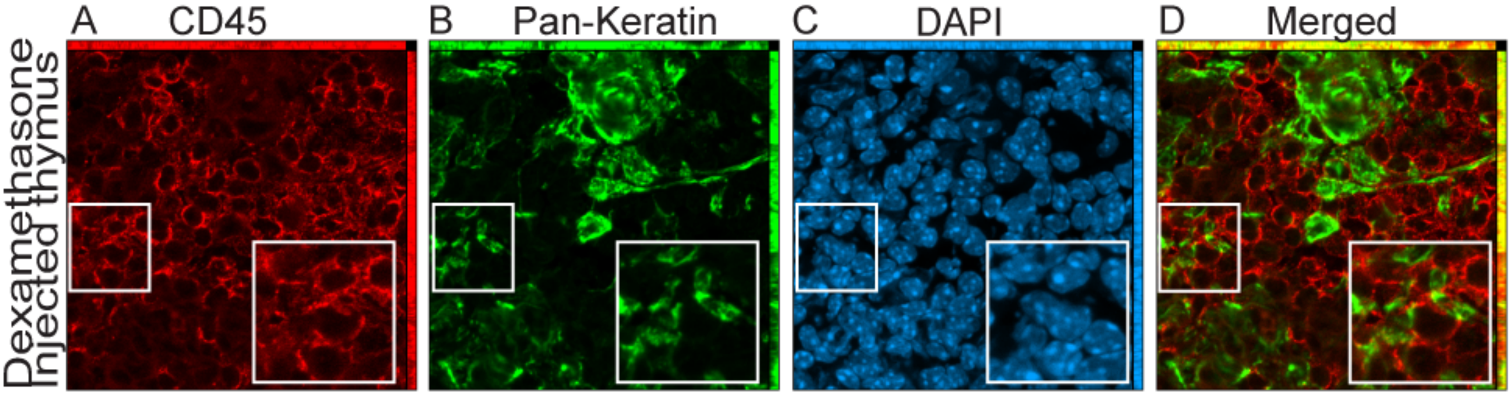
A
CD45
B
Pan-Keratin
C
DAPI
D
Merged
Dexamethasone
Injected thymus

S1. Presence of a CD45+Pank+ population in adult thymic sections. Dexamethasone treated murine thymuses were sectioned and stained for CD45 and Pan-Cytokeratin. Panel A showing a Maximum Intensity projection of CD45 staining; Panel B showing Maximum Intensity projection of Pan-Keratin staining; Panel C showing Maximum Intensity projection of DAPI staining. Panel D showing Maximum Intensity projection of merged CD45 and EpCAM staining. CD45 and EpCAM expressing cell of interest is enlarged in the insets to show co-expression of both proteins and represent the cell shown in the white boxes.

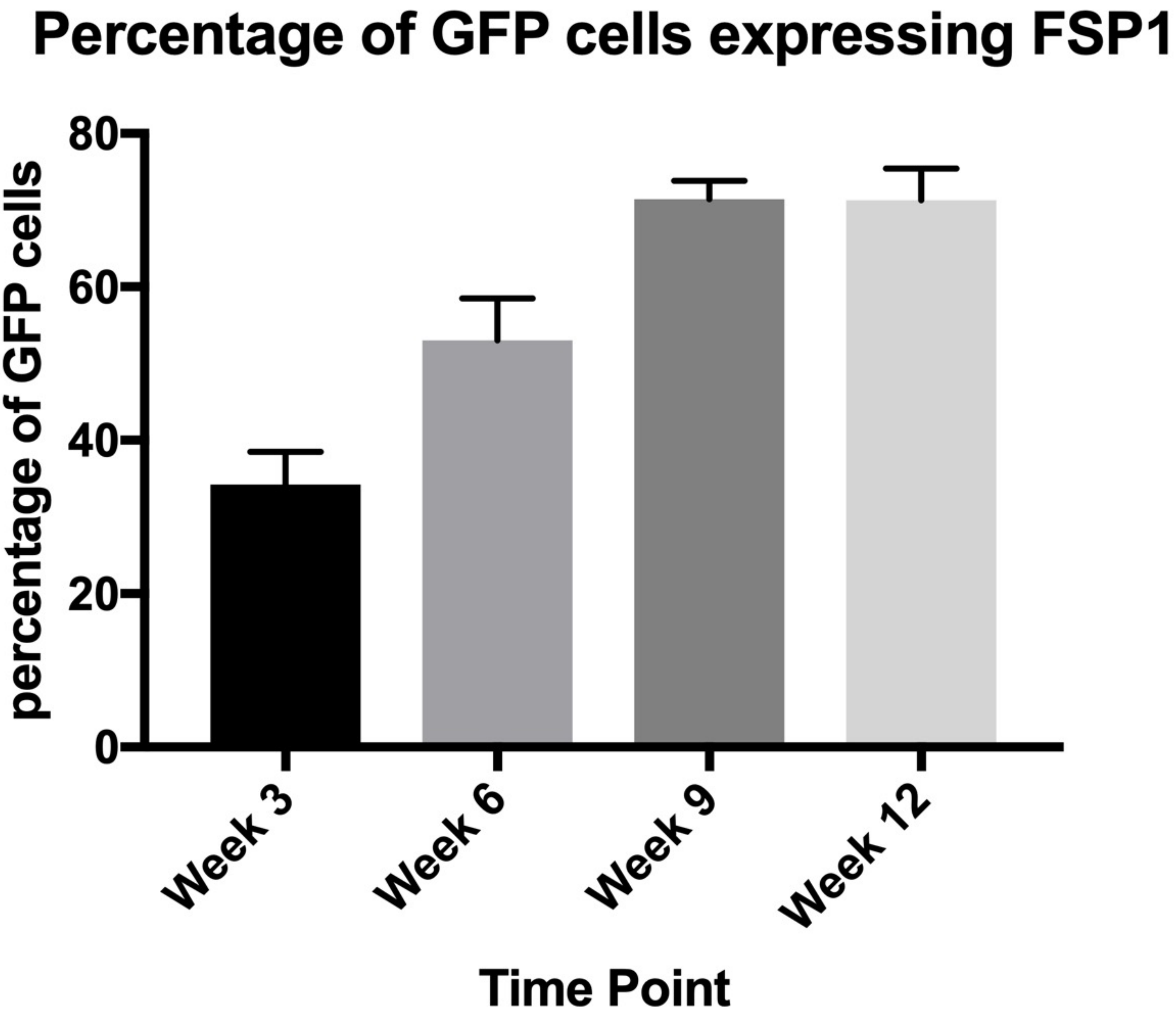
Percentage of GFP cells expressing FSP1
percentage of GFP cells
0
20
40
60
80
Week 3
Week 6
Week 9
Week 12
Time Point

S 2. Peripheral cells migrating into the thymus contribute to FSP1-expressing fibroblasts. C57BL6 fetal thymuses were transplanted under the kidney capsule of Actin H2BGFP mice and analyzed 3-12 weeks after transplant for the presence of GFP-expressing cells migrating into the transplanted lobes using IHC. Bar-Graph showing the mean of frequency of GFP expressing peripheral cells also expressing FSP1 at different timepoints following transplant (error bars represent SEM).

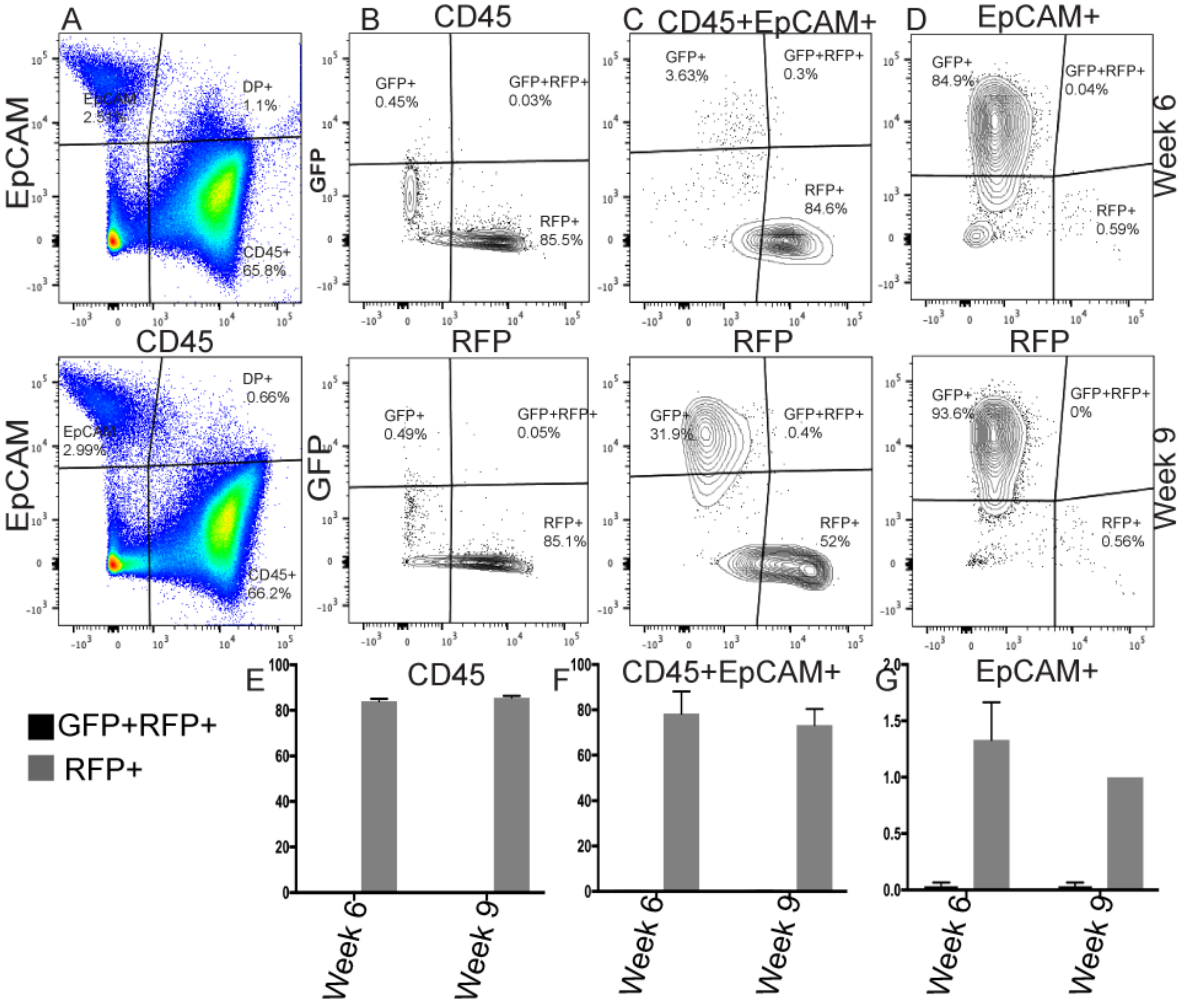

A
EpCAM
CD45
EpCAM
2.51%
DP+
1.1%
CD45+
65.8%
EpCAM
2.99%
DP+
0.66%
CD45+
66.2%
B
CD45
GFP
GFP+
0.45%
GFP+RFP+
0.03%
RFP+
85.5%
RFP
GFP+
0.49%
GFP+RFP+
0.05%
RFP+
85.1%
RFP
C
CD45+EpCAM+
GFP+
3.63%
GFP+RFP+
0.3%
RFP+
84.6%
RFP
GFP+
31.9%
GFP+RFP+
0.4%
RFP+
52%
RFP
D
EpCAM+
GFP+
84.9%
GFP+RFP+
0.04%
RFP+
0.59%
RFP
GFP+
93.6%
GFP+RFP+
0%
RFP+
0.56%
RFP
Week 6
Week 9
E
CD45
F
CD45+EpCAM+
G
EpCAM+
GFP+RFP+
RFP+
Week 6
Week 9

S 3. The CD45+EpCAM+ and CD45-EpCAM+ TEC populations derived from peripheral cells are not result of cell fusion. Actin H2B-GFP fetal thymuses were transplanted under the kidney capsule of mRFP ROSA26 mice and analyzed 6 and 9 weeks after transplant using FACS. All the experiments were done in triplicate. A: The FACS representation of the lineage depleted cells derived from dissociated engrafted thymi gated for CD45 and EpCAM B: The FACS representation of the percentage of GFP+; GFP+RFP+; and RFP+ expressing cells within the CD45+EpCAM- population at different timepoints; C: The FACS representation of the percentage of GFP+; GFP+RFP+; and RFP+ expressing cells within the CD45+EpCAM+ population at different timepoints; D: The FACS representation of the percentage of GFP+; GFP+RFP+; and RFP+ expressing cells within the CD45-EpCAM+ population at different timepoints. E. Bar graph showing the mean percentage of the CD45+GFP+RFP+ (potential cell fusions) and CD45+RFP+ (peripheral cells) populations at different time-points (error bars represent SEM). F: Bar graph showing the mean percentages of the CD45+EpCAM+GFP+RFP+ (potential cell fusions); CD45+EpCAM+RFP+ (peripheral cells) populations at different time-point (error bars represent SEM). G: Bar graph showing the percentages of the EpCAM+GFP+RFP+ (potential cell fusions); EpCAM+RFP+ (peripheral cells) cell populations at different time-points (error bars represent SEM).
